# PeakLocus: A transcript-aware toolkit for RNA-Centric Genomic Annotation

**DOI:** 10.64898/2026.10.17.759721

**Authors:** George Boateng-Sarfo, Shashank Pritam, Sarah Signor

## Abstract

Annotating genomic coordinates is an important step for identifying where along the transcript genomic features are enriched yet many widely used peak annotation tools were designed primarily around general genomic or chromatin features. We developed PeakLocus, a dependency-free Java command line tool that assigns RNA feature to transcript interval in an explicitly RNA centered transcript context. PeakLocus accepts bed-like and tabular interval files together with a gene transfer file as reference, detects coordinates and strand structure, and reports gene, transcript, biotype, and location information such as 5’ UTR, 3’ UTR, CDS. Generic UTR features are resolved using transcript-specific CDS boundaries and strand, while introns are derived from transcript exon structure. We benchmarked PeakLocus against BEDTools, UROPA, ChIPseeker, and HOMER using 466,526 evaluable RNA-associated peaks from 27 Drosophila, mouse, and human datasets and a balanced reference-derived benchmark of 30,000 fixed-width intervals. On real-world data, PeakLocus assigned a strict RNA-location category to 92.8% of evaluable intervals and showed 99.5% concordance with a direct BEDTools-derived overlap baseline among jointly annotated peaks. In the controlled benchmark, PeakLocus achieved 86.4% exact five-class recovery overall, with 99.9% recovery in mouse and 99.6% in human. Comparator differences were strongly influenced by native annotation ontology, including CDS-versus-exon resolution, intron representation, and non-RNA categories. PeakLocus provides a lightweight and interpretable framework for transcript-aware annotation of RNA-associated genomic intervals.

## Introduction

RNA high throughput assays produce genomic intervals rather than only gene-level measurements. Examples include N^6^-methyladenosine (m^6^A)^1^ and other RNA-modification peaks, Cross Linking and immunoprecipitation^2,3^ and related RNA-binding protein assays, direct RNA sequencing-derived modification sites^4^, and transcript-associated peak calls^5–9^. Interpreting these intervals requires more than assigning a nearby gene. The biological meaning of a site can depend on whether it falls in a 5’ untranslated region (5’ UTR), coding sequence (CDS), 3’ untranslated region (3’ UTR), other exonic sequence, or an intron^10–12^. The importance of transcript context is particularly clear for RNA modifications such as m^6^A, whose distribution is strongly structured across transcript regions and whose effects can depend on location^13,14^.

General-purpose genomic annotation tools provide powerful and flexible approaches for intersecting or assigning genomic ranges. BEDTools offers a low-level and highly reproducible interval-overlap framework^15^, UROPA supports configurable GTF-based genomic range annotation^16^, ChIPseeker provides peak annotation and visualization with a strong emphasis on ChIP-seq and nearest-gene/TSS relationships^16^, and HOMER provides broad genomic annotation as part of a larger regulatory genomics toolkit^17^. These tools, however, differ in their native feature vocabularies, prioritization rules, gene-assignment strategies, and treatment of transcript structure. Consequently, the same RNA-associated interval can receive different labels depending on whether a workflow emphasizes direct overlap, nearest gene, promoter/TSS context, or transcript features.

A second challenge is reference heterogeneity. Human and mouse GENCODE annotations use rich transcript models and commonly encode UTR sequence using a generic UTR feature, whereas FlyBase uses species-specific feature conventions^10,11^. Annotation software therefore needs to interpret reference structure rather than assume that all GTF files use identical feature labels. Noncoding transcripts create an additional challenge because exonic sequence should not be forced into CDS or 5’/3’ UTR categories when those assignments are not biologically defined.

PeakLocus was developed to provide a small, portable, RNA-centered annotation layer for these use cases. The program is implemented in Java, has no external Java dependencies, accepts common genomic interval formats, and returns an intentionally restricted RNA-location vocabulary: 3’ UTR, 5’ UTR, CDS, exon, intron, and N/A. The design goal is not to replace general-purpose regulatory annotation tools, but to answer a specific transcript-oriented question: which annotated RNA region does this genomic interval overlap?

Here, we describe the PeakLocus annotation model and evaluate it using two complementary benchmarking strategies. First, we compare PeakLocus with BEDTools, UROPA, ChIPseeker, and HOMER across 466,526 evaluable RNA-associated peaks from 27 public Drosophila, mouse, and human datasets obtained from GEO series GSE143821^18^, GSE192440^7^, and GSE114150^6^, respectively. Second, we construct a balanced, reference-derived benchmark containing 30,000 fixed-width intervals from five RNA-location classes across the same three species^10–12^. The combined analysis evaluates annotation coverage, pairwise location concordance, gene-ID concordance, exact five-class recovery, class-specific precision/recall/F1, and the effect of tool ontology on apparent disagreement.

## Materials and Methods

### Software implementation and input/output model

PeakLocus v1.0.0 was implemented in Java with Java 8 bytecode compatibility and no third-party Java dependencies. The program accepts plain-text or gzip-compressed genomic interval files, including BED3/BED4/BED5/BED6/BED12, narrowPeak, broadPeak, and custom tabular files. A GTF annotation file is supplied as the transcript reference. Each input row is preserved and PeakLocus appends Gene ID, Gene Name, Transcript IDs, Gene Type, Transcript Type, Gene Strand, and Location.

A typical invocation is:

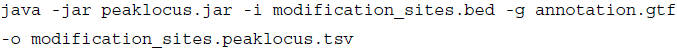

The program is distributed as a command-line JAR and can be incorporated into shell, SLURM, Snakemake, or other workflow systems. The source repository and documentation are available from the project repository listed in the Code Availability section.

## Mathematical representation of genomic intervals

### Interval representation

Let an experimental query interval be represented as

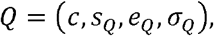

where *c* is the chromosome or contig, *S_Q_* and *e_q_* are the normalized inclusive start and end coordinates, and σ*Q* ɛ{+, −,} denotes positive strand, negative strand, or unknown/unstranded orientation.

A reference feature is represented as

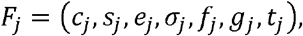

where *f_j_* is the GTF feature class, *g_j_* is the associated gene, and *t_j_* is the associated transcript. A reference feature can be considered for a query only when the chromosome matches:

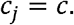

The strand search set is defined by the query strand as

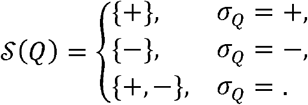

Thus, known strand restricts the search to concordant transcript annotations, whereas an unstranded interval is evaluated against both orientations.

### Coordinate normalization

GTF coordinates are 1-based and inclusive, while BED and MACS coordinate conventionally use 0-based half-open intervals, in which the start coordinate is included, and the end coordinate is excluded.

For BED-like input, PeakLocus converts the query internally as

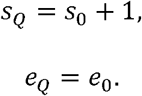

Therefore, PeakLocus applies the BED-to-GTF normalization rule:

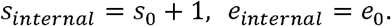

For input already specified as 1-based inclusive,

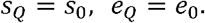

Only the internal query coordinates are transformed. PeakLocus preserves the original input coordinates in the output file.

#### 2.2.3 Base-pair overlap

For a normalized query interval *Q* = [*s_Q_*, *e_Q_*] and a reference feature *F* = [*s_F_*, *e_F_*], the number of overlapping bases is

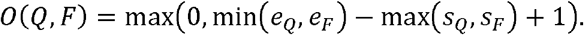

An overlap exists if and only if

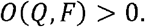

The overlap length is subsequently used both to establish intersection and to break ties between competing annotations of the same RNA-region priority.

### Strand handling and automatic strand-column detection

#### Motivation

RNA-associated interval files do not universally encode strand in BED column 6. For example, a modkit-derived table may place strand in an earlier column, while custom tables can place orientation almost anywhere. PeakLocus therefore detects strand automatically rather than assuming a fixed field.

By default, the detector samples up to

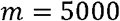

input rows during schema inference.

#### Strand token normalization

PeakLocus maps the following informative tokens to a positive or negative orientation:

### Strand normalization

PeakLocus standardizes heterogeneous strand encodings into positive (+), negative (-), or unstranded states. Positive-strand synonyms include plus, forward, fwd, and sense, whereas negative-strand synonyms include minus, reverse, rev, antisense, and anti-sense. Missing or non-informative values, including., *, NA, N/A, none, unknown, and unstranded, are treated as unstranded. Unstranded intervals are retained and queried without imposing a strand restriction.

Column-wise strand profile

For a candidate column *c*, let

- *n_c_* = number of observed non-empty values,
- *r_c_* = number of values recognized as either informative strand or accepted unstranded tokens,
- *i_c_* = number of informative positive or negative strand values.

The recognized fraction is

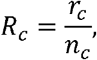

and the informative fraction is

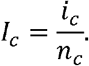

For arbitrary value-profile detection, PeakLocus requires

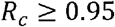

and

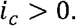

The strand-likeness score is

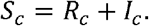

Among qualifying arbitrary columns, the selected strand column is

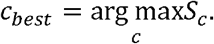

The requirement for informative + or − evidence prevents columns containing only symbols such as., NA, or * from being misidentified as strand fields.

#### 2.3.4 Detection hierarchy

PeakLocus applies strand detection in a predefined hierarchy.

Step 1: header-based detection

Header names equal to or containing strand receive priority. orientation and sense are treated as weaker header signals. For a candidate header column, the implementation calculates a score of the form

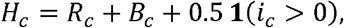

where *B_c_* = 2 for strong strand-associated header names and *B_c_* = 0 for weak orientation labels. The indicator function **1**(*i_c_* > 0) rewards direct +/− evidence.

Step 2: BED6/MACS convention

If the input has at least six columns, column 6 is recognized as a conventional BED/MACS strand field when

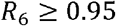

and at least one recognized informative or unstranded token is present.

Step 3: arbitrary-column profiling

If neither header detection nor the BED6 convention resolves strand, all columns are profiled using *S_c_* = *R_c_* + *I_c_*, subject to the requirements above.

Step 4: conservative unstranded fallback

If no reliable strand column is identified,

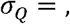

and the annotation engine queries both transcript strands. PeakLocus therefore avoids inventing orientation when the input does not support it.

### GTF preprocessing and interval indexing

#### RNA-relevant feature extraction

PeakLocus indexes transcript-relevant GTF features including exon, CDS, generic UTR, explicit five-prime and three-prime UTR labels, start codon and stop codon. Transcript records are retained as metadata used during intron construction but are not treated as generic gene-body overlaps.

Gene and transcript metadata are recovered from GTF attributes. PeakLocus accepts either gene_type or gene_biotype, and either transcript_type or transcript_biotype.

#### 2.4.2 Transcript-specific CDS boundaries

For transcript *t*, define the set of CDS features as *CDS*(*t*). PeakLocus calculates

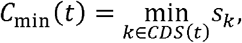

and

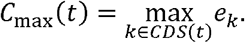

These transcript-specific bounds are used to orient generic GENCODE UTR records.

##### Sorted interval index

Reference records are partitioned by chromosome and strand and then sorted by genomic start coordinate:

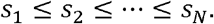

For every position *i*, PeakLocus stores a prefix maximum end coordinate

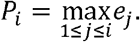

This array permits safe early termination during backward interval scanning.

##### Query boundary by binary search

For query *Q* = [*q_s_*, *q_e_*], binary search identifies the first sorted reference position *p* satisfying

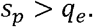

Only records with index less than *p* can begin early enough to overlap the query. PeakLocus scans backward from *p* – 1.

The scan terminates when

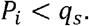

Because

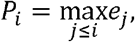

the condition *P_i_* < *q_s_* implies

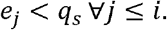

Therefore, no earlier indexed feature can overlap the query. This stopping rule avoids the failure mode in which a long gene or transcript begins well before a query but extends across it while shorter neighboring records end earlier.

### Transcript-aware RNA feature construction

#### Generic UTR inference

If a GTF explicitly labels a feature as five_prime_utr or three_prime_utr, PeakLocus accepts that label directly. For a generic UTR, the feature is classified relative to the transcript-specific CDS boundaries.

Let a generic UTR interval be *U* = [*S_U_*, *e_U_*].

For a positive-strand transcript,

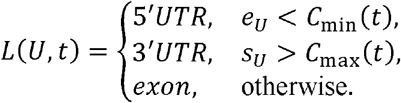

For a negative-strand transcript, transcriptional orientation is reversed:

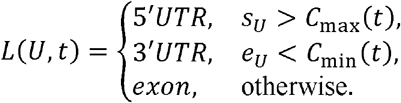

If no usable CDS bounds exist for transcript *t*, PeakLocus conservatively assigns

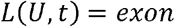

rather than inventing a 5’/3’ orientation. The same conservative exon fallback is used for rare inconsistent generic UTR annotations that overlap the inferred CDS bounds.

### Transcript-aware intron construction

PeakLocus constructs introns from gaps between consecutive exon blocks for each transcript independently.

For transcript *t*, let the exons sorted by genomic start be

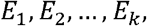

where

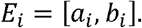

Because annotated exons can overlap, PeakLocus maintains a merged exon endpoint. Initialize

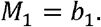

For each subsequent exon *E_i_*, an intron is created only if

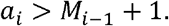

The intron interval is then

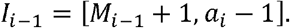

After evaluating the gap, the merged endpoint becomes

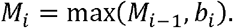

This construction prevents overlapping exon annotations from creating false introns and retains transcript specificity in alternatively spliced genes.

### RNA-region classification and annotation priority

#### Structural classification

Each overlapping GTF-derived record is converted to one of the allowed RNA-region categories. Explicit 5’ UTR and 3’ UTR records are retained, CDS and Selenocysteine are classified as CDS, exons as exon, and reconstructed transcript gaps as intron. Consistent with the precursor algorithm, start-codon records are associated with 5’ UTR and stop-codon records with 3’ UTR.

#### 2.6.2 Annotation priority

When one query interval overlaps multiple RNA features, PeakLocus first differentiate between more specific subregions and broader structural annotations. The RNA class are assigned a specific rank

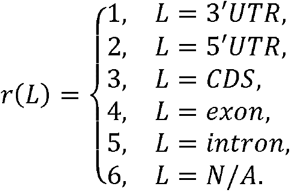

Thus, for each overlapping candidate, hi, PeakLocus also calculates the number of overlapping bases

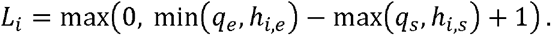

candidate annotation are them compared using. Hence, PeakLocus, first retains the most specific compatible RNA region annotation and, when multiple candidate have the same annotationm rank, selects the feature with the largest base-pair overlap.

#### 2.6.3 Deterministic selection of the best hit

Let the set of overlapping candidate hits for query *Q* be

Each hit has priority *π*(*h*), overlap length *O*(*h*), gene identifier *G*(*h*), and transcript identifier *T*(*h*). PeakLocus orders hits lexicographically by

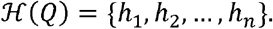

The selected hit is

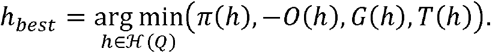

This implements the following rules:

1. Retain the most specific compactible RNA region annotation;
2. among equal-priority hits, prefer the largest base-pair overlap;
3. use gene identifier and transcript identifier as deterministic tie-breakers.

After selecting *h_best_*, PeakLocus retains the set of transcript identifiers that match the selected gene, strand, priority, and location:

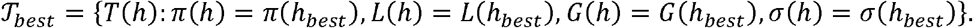

The resulting transcript identifiers are written as a semicolon-delimited list.

**Table 1.** PeakLocus RNA-location classification framework.

| Priority | Location | Interpretation |
| --- | --- | --- |
| 1 | 3' UTR | Annotated or transcript inferred three-prime untranslated region |
| 2 | 5' UTR | Annotated or transcript five-prime untranslated region |
| 3 | CDS | Coding sequence |
| 4 | exon | Exonic sequence not resolved to UTR or CDS |
| 5 | intron | Transcript-aware intronic sequence |
| 6 | N/A | No compatible RNA-location assignment |

### 2.7 Annotation of coding and noncoding RNA biotypes

PeakLocus does not filter reference records to protein-coding genes. Gene and transcript biotypes are carried directly from the GTF annotation. Consequently, PeakLocus can preserve annotations for any RNA class represented in the supplied GTF, including protein-coding transcripts, lncRNAs, snRNAs, snoRNAs, miRNAs, rRNAs, and other noncoding or processed transcript classes.

RNA identity and RNA structural position are treated as separate attributes. For example, an snRNA may be represented as Gene_Type = snRNA, Transcript_Type = snRNA, Location = exon. Whereas a protein-coding transcript may be represented as Gene_Type = protein_coding, Transcript_Type = protein_coding, Location = 3’ UTR. This separation avoids incorrectly using an RNA biotype as a structural location category. For noncoding RNAs without CDS-based UTR structure, exon and transcript-derived intron remain valid structural annotations.

### 2.8 Computational complexity

Let *N* be the number of indexed reference features and *M* the number of query intervals.

Sorting the reference intervals requires approximately

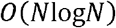

time. Construction of the prefix maximum-end arrays requires

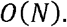

Exons must also be grouped and sorted by transcript to construct transcript-aware introns. If *N_E_* is the number of exon records, an upper-bound description of this preprocessing step is

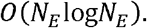

For a query against a chromosome/strand partition containing *N_c_* indexed records, the candidate boundary is identified by binary search in

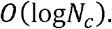

If *K* records are inspected during the subsequent backward scan before the prefix maximum-end condition terminates the search, the per-query complexity is approximately

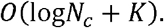

The worst case remains linear in an unusually dense or deeply overlapping partition, but the prefix maximum-end rule avoids a complete scan for typical localized queries.

The reference index, metadata, CDS bounds, exon groups, and derived introns require memory proportional to the annotation size, approximately

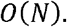

Input schema and strand detection samples at most *m* = 5000 rows across *C* columns, yielding

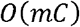

schema-detection work.

### 2.9 Software validation and self-tests

PeakLocus v1.0.0 includes dependency-free Java self-tests and an end-to-end command-line integration test. The current synthetic test suite evaluates:

1. detection of a modkit-style strand field outside BED column 6;
2. standard BED6 strand detection;
3. custom header-based strand detection;
4. headerless arbitrary-column strand detection;
5. unstranded BED3 fallback;
6. positive-strand 5’ UTR, CDS, and 3’ UTR assignment;
7. negative-strand reversal of 5’/3’ UTR interpretation;
8. transcript-aware intron construction;
9. noncoding exon annotation; and
10. complete command-line output generation.

These tests establish algorithmic correctness against synthetic examples for which the expected annotation is known. They should not be interpreted as a substitute for large-scale biological benchmarking.

### 2.10 Reference annotations, comparator tools, and harmonization

The benchmark used species-specific GTF references matching the processed datasets: FlyBase Drosophila melanogaster release 6.68 (dmel-all-r6.68.gtf), GENCODE mouse vM23, and GENCODE human v50. GENCODE provides curated human and mouse gene/transcript annotations^10^, while FlyBase provides *Drosophila* gene and genome annotations^11^. Because the feature vocabularies differ among these references, species-specific native feature names were retained during tool execution and harmonized only during comparative scoring.

#### 2.10.1 Comparator tools and execution

PeakLocus was benchmarked against BEDTools v2.31.1 ^15^, UROPA v4.0.3 ^16^, ChIPseeker v1.48.0^19^, and HOMER v5.1^17^. Comparator tools were executed with the same species-specific GTF used by PeakLocus wherever supported. BEDTools intersect was used as a direct overlap baseline with all overlapping GTF records retained. UROPA was run at zero distance with internal overlaps enabled and native RNA-relevant feature names appropriate to each GTF. ChIPseeker used transcript-level annotation with a –3 kb/+3 kb TSS window and its native annotation categories. HOMER annotatePeaks.pl was run in annotation-only mode using the supplied GTF.

**Table 2.** Comparator configuration used in the benchmark.

| Tool | Version | Primary benchmark behavior |
| --- | --- | --- |
| PeakLocus | 1.0.0 | Transcript-aware RNA-location annotation; priority 3' UTR > 5' UTR > CDS > exon > intron > N/A |
| BEDTools | 2.31.1 | All direct GTF overlaps retained;<br>RNA class derived during<br>harmonization |
| UROPA | 4.0.3 | Zero-distance internal overlap<br>using native feature vocabulary |
| ChIPseeker | 1.48.0 | Transcript-level annotatePeak;<br>native genomic feature categories |
| HOMER | 5.1 | annotatePeaks.pl with supplied<br>GTF |

#### 2.10.2 Strict cross-tool harmonization

Native tool outputs were retained and then harmonized into 3’ UTR, 5’ UTR, CDS, exon, intron, and N/A. Only transcript-relevant categories counted as strict RNA annotations. Promoter, promoter-TSS, TTS, downstream, distal intergenic, intergenic, and other regulatory categories were retained in native audits but mapped to N/A for strict RNA-location comparisons. We did not reinterpret promoter as 5’ UTR or TTS as 3’ UTR. Generic GENCODE UTR records were oriented using transcript/CDS structure.

For BEDTools, raw overlaps were converted into an RNA-location prediction using the same feature-priority logic used for direct reference overlap. Gene/transcript-span evidence without a higher-priority exonic, UTR, or CDS feature was treated as intronic in this derived baseline. Consequently, the BEDTools-derived benchmark should be interpreted as a reference-overlap positive control rather than a fully independent annotation algorithm.

##### Real-world benchmark

Public RNA-associated peak datasets were selected from three species and three independent studies. The *Drosophila* benchmark used six m^6^A peak datasets from GSE143821, generated in the study of m^6^A-dependent FMR1 granule dynamics during early embryogenesis^18^. The mouse benchmark used twelve IP peak datasets covering GV oocyte, MII oocyte, zygote, 2-cell, 8-cell, and blastocyst stages from GSE192440. The human benchmark used nine processed peak files from GSE114150, which profiled m^6^A across fetal tissues.

The combined input contained 466,529 interval rows. Three human intervals on KI270733.1 were not evaluable because that contig was absent from the GENCODE v50 GTF used for comparative annotation. These records were retained in source files but excluded from comparative metrics, leaving 466,526 evaluable intervals. Duplicate genomic coordinates were not collapsed; each input row was tracked with a stable row identifier, so that replicated or repeated peaks remained independent benchmark observations.

**Table 3.**
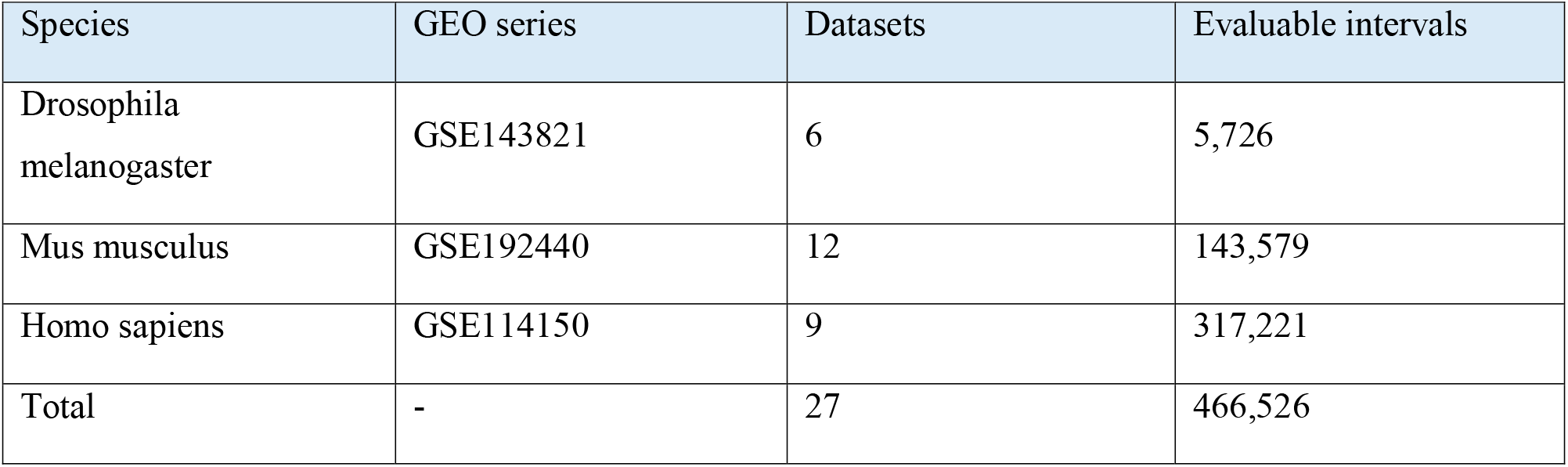
Real-world benchmark composition.

##### Real-world benchmark metrics

Strict RNA-feature coverage was defined as the proportion of evaluable intervals assigned to 3’ UTR, 5’ UTR, CDS, exon, or intron rather than N/A. Pairwise RNA-location concordance was calculated among intervals for which both tools assigned a non-N/A RNA-location category. This metric therefore quantifies exact category agreement among jointly annotated peaks and is not a global accuracy estimate.

Gene-ID concordance with PeakLocus was calculated only where both PeakLocus and the comparator reported at least one gene identifier. A peak was considered concordant when the reported gene-ID sets shared at least one gene. Because BEDTools/PeakLocus emphasize direct overlap while UROPA, ChIPseeker, and HOMER can use feature-associated or nearest-gene semantics, gene concordance was treated as a descriptive comparison rather than gene-assignment accuracy.

### Controlled reference-derived truth benchmark

A balanced controlled benchmark was generated from the same species-specific GTF references. Transcript structures were parsed to obtain exons, CDS segments, UTRs, and transcript-aware introns. GTF 1-based inclusive coordinates were converted to BED-style 0-based half-open coordinates. Generic GENCODE UTRs were separated into 5’ UTR and 3’ UTR using transcript strand and CDS boundaries.

Reference features were decomposed into atomic genomic segments. Primary truth masks retained only segments associated with one active gene, one strand, and one RNA-location class, thereby excluding overlapping-gene and isoform-discordant regions. Adjacent compatible atomic segments were merged. One 100-bp interval was then sampled from each distinct eligible source region using a fixed random seed (20260926). For each species, 2,000 intervals were sampled from each of the five classes (3’ UTR, 5’ UTR, CDS, exon, intron), producing 10,000 intervals per species and 30,000 intervals overall.

All five tools were run on the same 30,000 intervals, yielding 150,000 tool-specific predictions. Controlled metrics included exact five-class recovery, balanced accuracy, class-specific precision/recall/F1, macro-F1, N/A rate, and gene-ID recovery where comparable identifiers were available. Because the truth labels were generated from the same GTFs supplied to the annotation tools, these measurements represent reference-consistent recovery rather than independent biological ground truth. In particular, the BEDTools-derived result serves as an internal positive control.

### Benchmark metrics and statistical definitions

For the controlled reference-derived benchmark, exact recovery was calculated by comparing each predicted RNA-location class with its reference-derived truth label. If *N_correct_* of *N_total_* intervals match the expected structural annotation, overall accuracy is

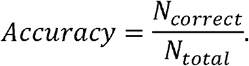

For each RNA-region class *r*, define true positives *TP_r_*, false positives *FP_r_*, and false negatives *FN_r_* T.

Then

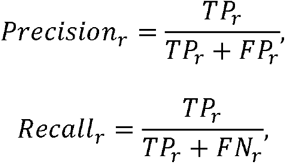

and

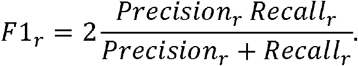

Class-specific confusion matrices were used to identify systematic substitutions among UTR, CDS, exon, and intron categories.

Strict RNA-feature coverage was calculated as:

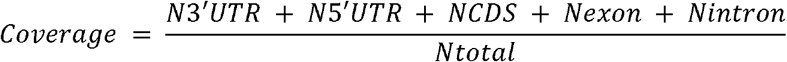

For two tools A and B, pairwise RNA-location concordance among jointly annotated non-N/A intervals was defined as:

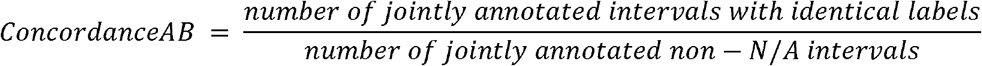

Macro-F1 was the unweighted mean of the five class-specific F1 scores:

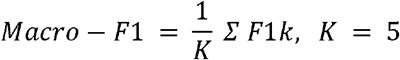

Balanced accuracy was the unweighted mean of class-specific recall:

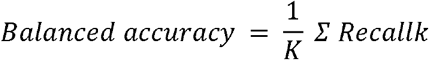

The fraction of intervals that could not be mapped to one of the five strict RNA classes was reported as the N/A rate:

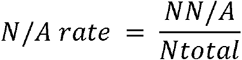

Because the controlled benchmark contained the same number of truth intervals in each of the five classes, overall accuracy and balanced accuracy are numerically identical for that benchmark. The benchmark statistics are deterministic summaries of the complete fixed test sets; no null-hypothesis significance test was used to reinterpret these fixed annotation outcomes.

## Results

### Benchmark design and RNA-annotation behavior

The real-world benchmark contained 466,526 evaluable RNA-associated intervals across 27 datasets: 5,726 Drosophila peaks from six datasets, 143,579 mouse peaks from twelve datasets, and 317,221 human peaks from nine datasets (Figure 1A). Three human intervals on KI270733.1 were excluded from comparative scoring because that contig was absent from the GENCODE v50 GTF used for annotation.

**Figure 1.**
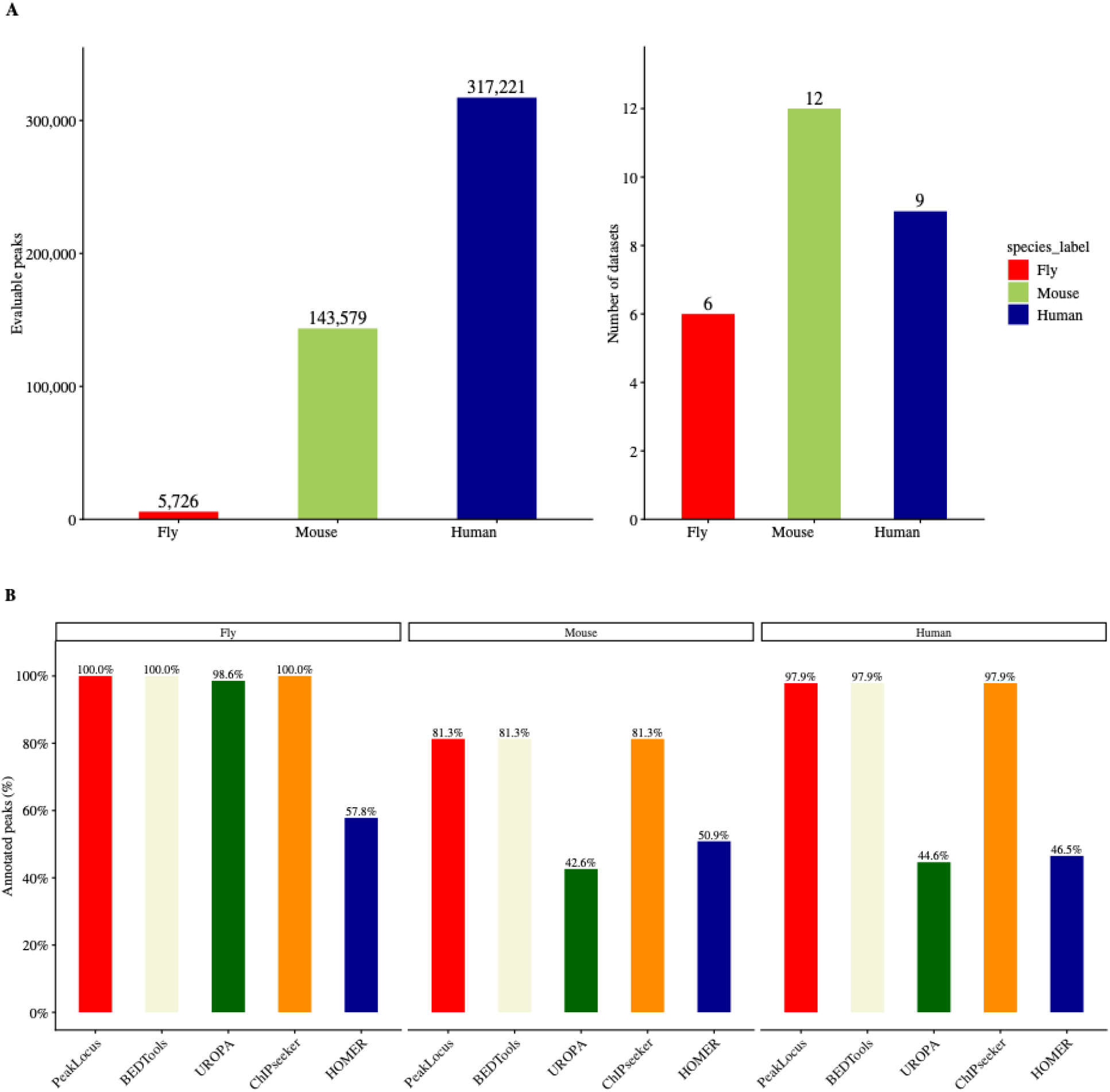
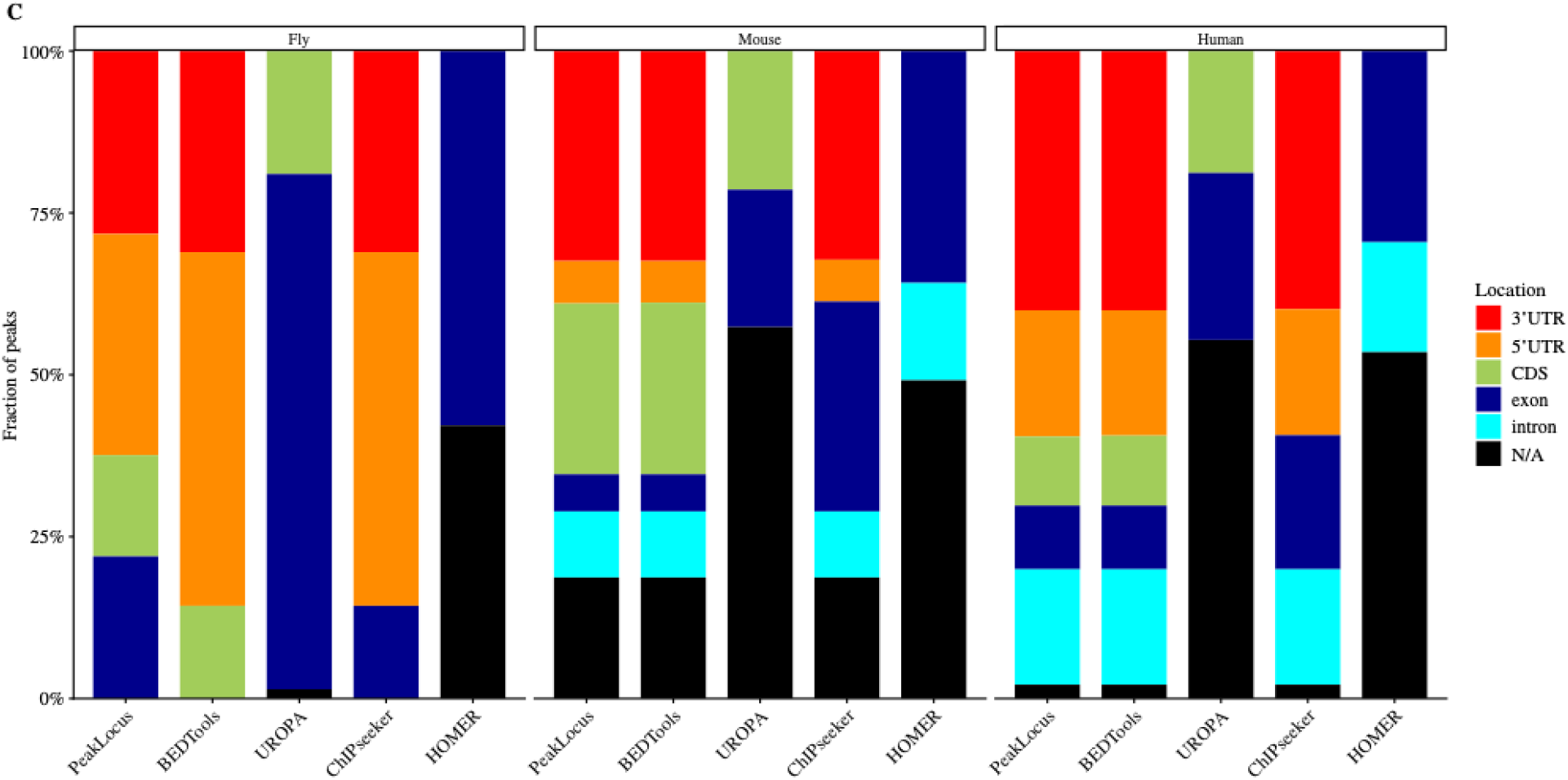
Benchmark design and RNA-annotation behavior. (A) Composition of the real-world benchmark across fly, mouse, and human. (B) Species-specific strict RNA-feature annotation coverage, where only 3’ UTR, 5’ UTR, CDS, exon, and intron count as strict RNA annotations. (C) Distribution of harmonized RNA-location categories across tools and species.

Strict RNA-feature coverage differed among tools and species (Figure 1B). PeakLocus, the BEDTools-derived overlap baseline, and ChIPseeker each annotated essentially all fly intervals and approximately 81-98% of mouse and human intervals with one of the five strict RNA classes. UROPA and HOMER produced lower strict RNA-feature coverage in mouse and human. Across all evaluable intervals, PeakLocus, BEDTools-derived, and ChIPseeker each reached approximately 92.8% strict RNA-feature coverage, compared with 44.7% for UROPA and 48.0% for HOMER.

The harmonized category distributions further showed that the tools differed not only in the number of intervals annotated but also in the type of RNA context reported (Figure 1C). PeakLocus and BEDTools-derived showed closely matched RNA-location profiles, while the comparator distributions reflected differences in native annotation vocabulary and feature-selection behavior.

#### Cross-tool RNA-location concordance

Among peaks for which both tools assigned a non-N/A strict RNA-location category, PeakLocus showed 99.5% exact concordance with the BEDTools-derived baseline (Figure 2A). Concordance with ChIPseeker was 82.6%, whereas concordance with UROPA and HOMER was 30.9% and 34.8%, respectively. The corresponding BEDTools comparisons were similar, with 82.9% agreement with ChIPseeker, 30.4% with UROPA, and 34.6% with HOMER.

**Figure 2.**
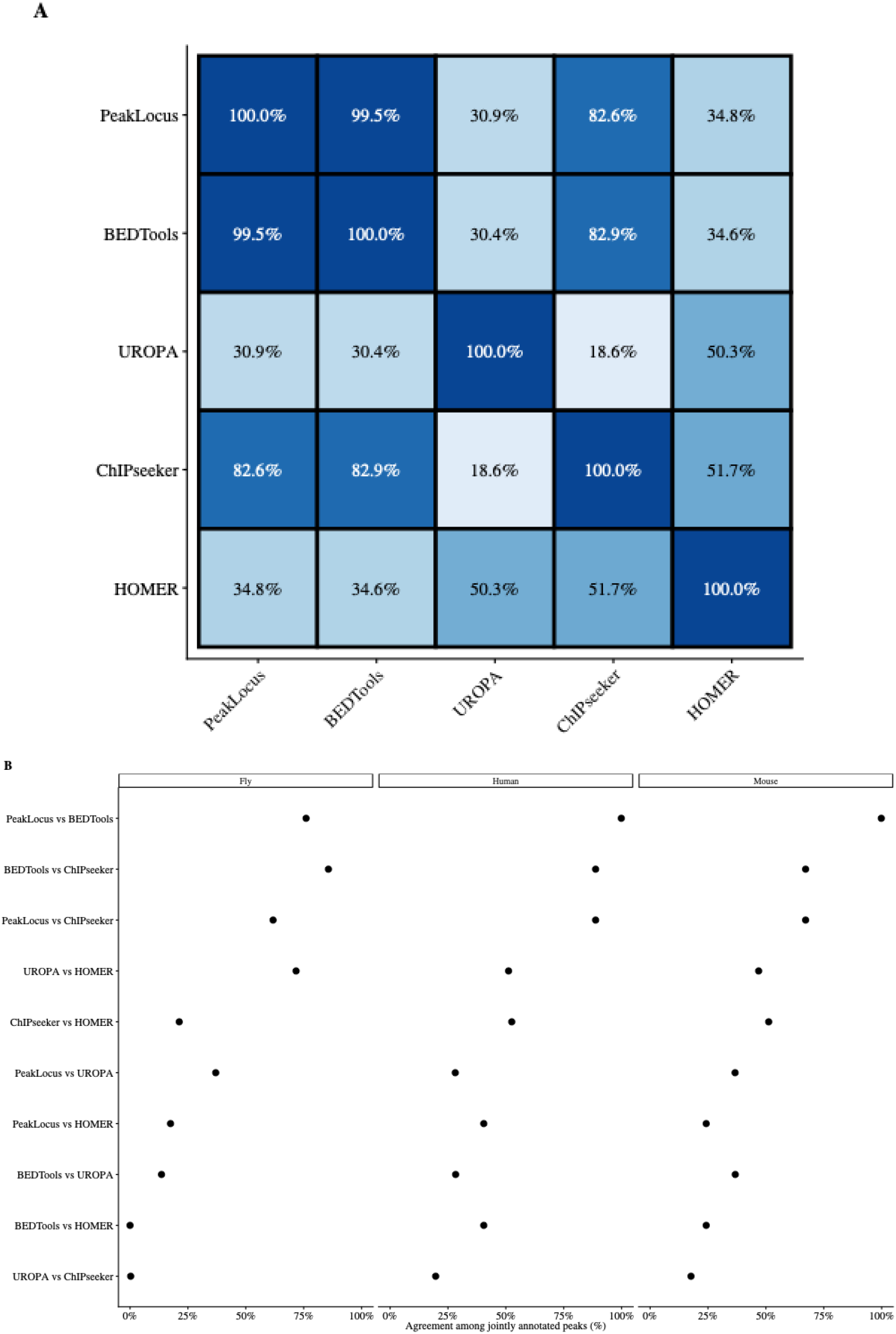
Cross-tool strict RNA-location concordance. (A) Pairwise exact category agreement among peaks for which both tools assigned a non-N/A RNA location. (B) The same pairwise concordance resolved by species.

Species-resolved comparisons retained the same general structure (Figure 2B): overlap-derived annotations remained the most concordant, while comparisons involving UROPA and HOMER showed substantially more divergence. These values measure agreement among jointly annotated peaks and should not be interpreted as independent biological accuracy.

#### Disagreement structure reveals differences in annotation resolution

Confusion-style comparison of PeakLocus with each comparator showed that disagreements were concentrated in specific category substitutions rather than distributed uniformly across the RNA-location vocabulary (Figure 3). PeakLocus and BEDTools-derived assignments were nearly identical for the dominant categories. ChIPseeker disagreements were enriched for differences between CDS and exon or among neighboring transcript-region labels, whereas UROPA and HOMER more often assigned broader or non-RNA-specific categories that became N/A after strict harmonization.

**Figure 3.**
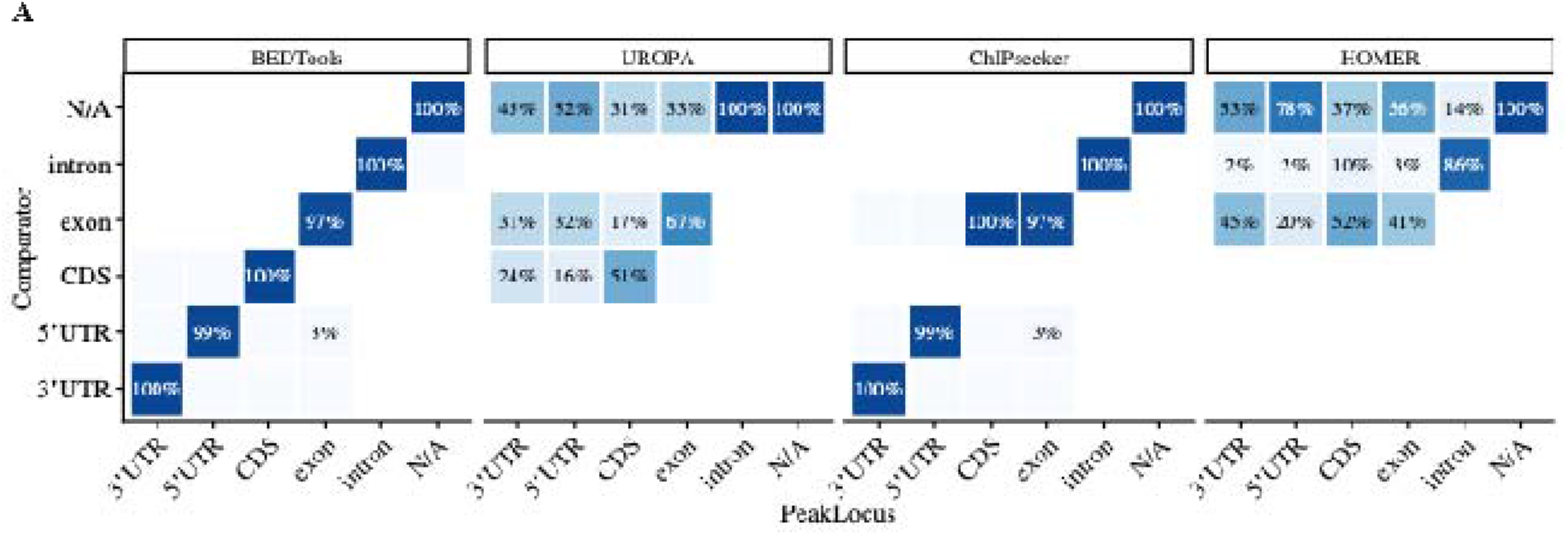
Structure of PeakLocus-comparator RNA-location disagreements. Cells show the percentage of each PeakLocus category assigned to each comparator category after strict harmonization.

#### Dataset-level robustness and reported gene-ID concordance

Dataset-level analysis showed that the species-level coverage patterns were reproduced across individual experimental datasets rather than being driven by a small number of samples (Figure 4A). PeakLocus, BEDTools-derived, and ChIPseeker remained consistently high within species, whereas UROPA and HOMER showed lower coverage and greater dataset-to-dataset variation.

**Figure 4.**
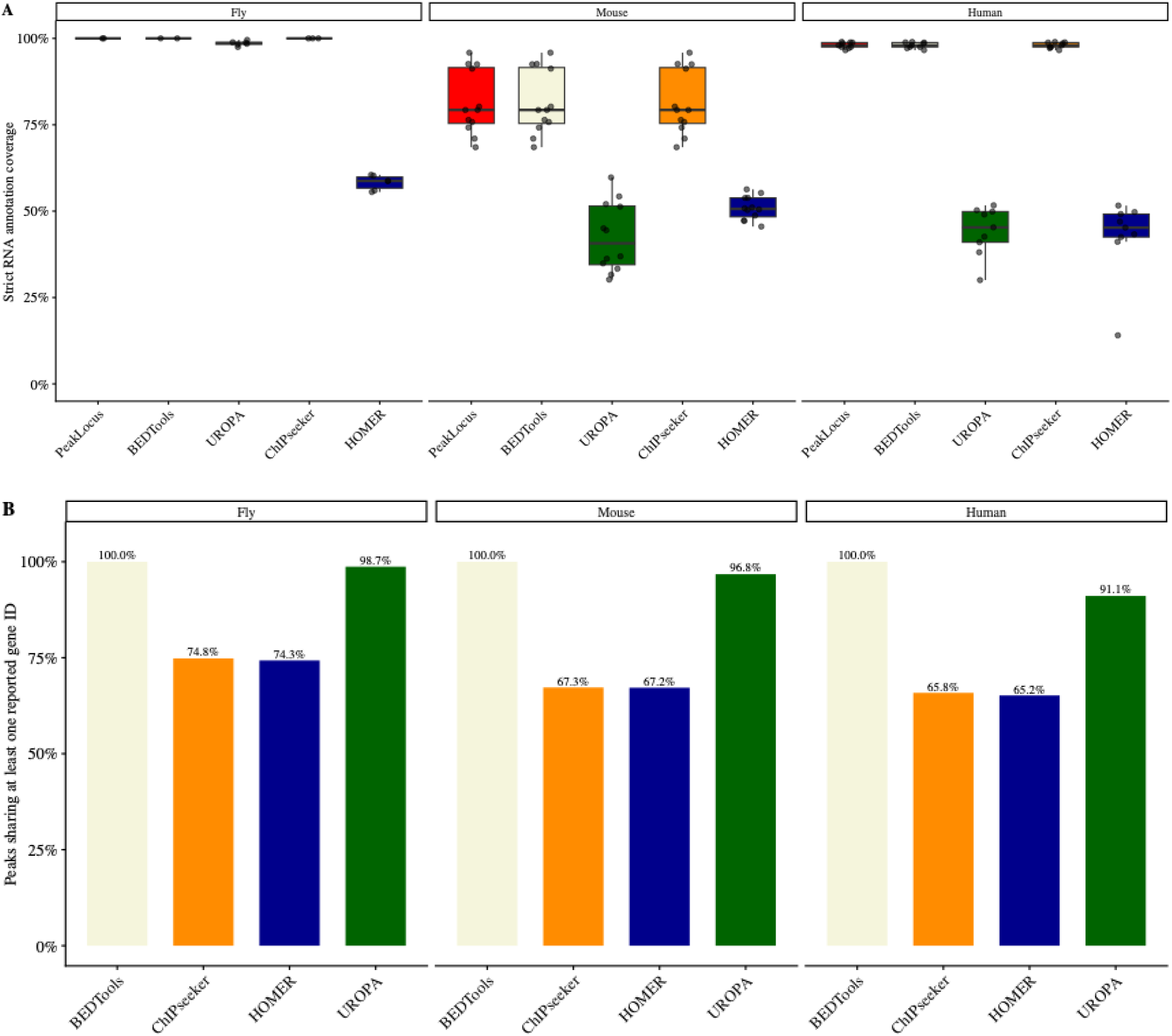
Robustness across datasets and reported gene-ID concordance. (A) Dataset-level strict RNA annotation coverage; each point represents one experimental dataset. (B) Fraction of evaluable peaks sharing at least one reported gene ID between PeakLocus and each comparator.

Reported gene-ID concordance with PeakLocus was 100% for the BEDTools-derived baseline in fly, mouse, and human (Figure 4B). UROPA shared at least one reported gene ID with PeakLocus for 98.7% of evaluable fly peaks, 96.8% of mouse peaks, and 91.1% of human peaks. ChIPseeker concordance was 74.8%, 67.3%, and 65.8%, respectively, while HOMER concordance was 74.3%, 67.2%, and 65.2%. Because these tools use overlapping-gene, feature-associated-gene, or nearest-gene assignment strategies, these values are interpreted as reported gene-ID concordance rather than gene-assignment accuracy.

### Controlled reference-derived benchmark

The controlled benchmark comprised 30,000 balanced 100-bp intervals, with 2,000 intervals from each of five RNA-location classes per species and produced 150,000 tool-specific predictions. All tools returned matched output for all 10,000 intervals in each species. Across all species, the BEDTools-derived reference-consistency baseline recovered 100.0% of the five-class labels, PeakLocus recovered 86.42%, ChIPseeker 79.99%, UROPA 69.15%, and HOMER 23.33% (Figure 5A). Macro-F1 values were 1.000, 0.868, 0.733, 0.713, and 0.233, respectively.

**Figure 5.**
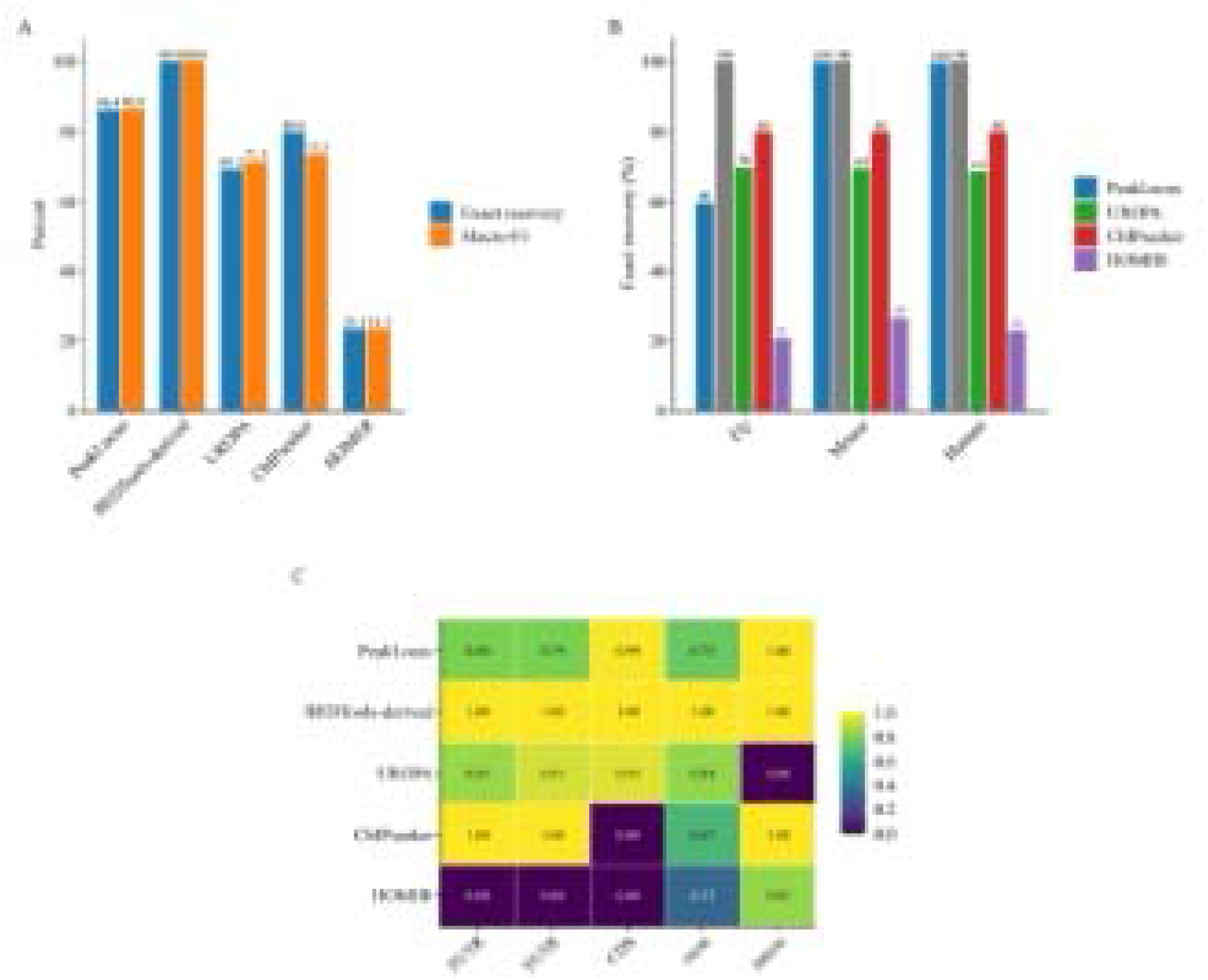
Controlled reference-derived benchmark. (A) Overall exact five-class recovery and macro-F1 across all species. (B) Species-specific exact recovery for each tool. (C) Per-class F1 across all species. BEDTools-derived is a reference-consistency positive-control baseline because the same GTF feature logic was used to derive the truth masks.

PeakLocus exact recovery was 99.87% in mouse and 99.64% in human, while Drosophila recovery was 59.76% in the exact-label benchmark (Figure 5B). The fly result was driven primarily by differences in the representation of UTR features under the FlyBase reference, while CDS, exon, and intron remained highly reproducible. In mouse and human, the remaining PeakLocus discrepancies were limited to a small number of coding/UTR boundary assignments under the benchmark’s strict class definition.

Per-class F1 scores exposed the native ontology limits of the comparator tools (Figure 5C). ChIPseeker recovered 3’ UTR and 5’ UTR with F1=1.00 and intron with approximately 1.00 but did not preserve CDS as a separate class in this workflow, causing CDS intervals to be represented within exon. UROPA showed strong F1 for 3’ UTR, 5’ UTR, CDS, and exon but no exact intron recovery under the strict harmonization because introns were not explicit GTF features and generic gene/transcript calls were not reinterpreted as intron. HOMER showed the lowest compatibility with the five-class RNA ontology and a 51.26% N/A rate.

## Discussion

PeakLocus was developed around a narrow but recurring bioinformatics problem which is that, translating genomic coordinates from RNA-associated experiments into transcript-region context. The software intentionally avoids promoter, enhancer, chromatin-state, and nearest-TSS labels because those concepts answer different biological questions. For RNA modifications, CLIP peaks, and direct-RNA-derived sites, distinguishing 5’ UTR, CDS, 3’ UTR, other exon, and intron can be more directly interpretable than assigning the closest transcription start site.

The real-world benchmark supports this design choice. PeakLocus assigned a strict RNA-location category to 92.8% of more than 466,000 evaluable peaks and closely reproduced the BEDTools-derived overlap baseline. At the same time, substantially lower concordance with UROPA and HOMER was accompanied by broader native feature vocabularies and larger N/A fractions after strict harmonization. The results therefore emphasize that annotation output is shaped by the biological ontology implemented by the tool, not only by coordinate overlap.

A central feature of the PeakLocus ontology is that exon is not synonymous with every sequence physically located in an exon. UTRs and CDS are themselves exonic but are reported using their more informative higher-priority labels. The exon category is reserved for sequence that is explicitly exonic but cannot be assigned to 3’ UTR, 5’ UTR, or CDS. This is particularly important for lncRNA and other noncoding transcripts, where a coding label would be inappropriate and a 5’/3’ UTR label may be undefined.

The controlled benchmark illustrates why this distinction matters. ChIPseeker achieved high localization of many transcript regions but did not preserve CDS as a separate class in the tested workflow, causing CDS truth intervals to appear as exon. This is not equivalent to a random localization failure; it is a loss of annotation resolution. Exact-category benchmarks should therefore be interpreted alongside the native ontology of each comparator.

### Direct-overlap baselines and independent comparators answer different questions

The BEDTools-derived baseline showed 99.5% real-world concordance with PeakLocus and 100% exact recovery in the controlled reference-derived benchmark. These results are expected because the baseline directly reconstructs RNA-location classes from the same reference features used by PeakLocus and by the truth-mask generator. It is therefore most useful as an implementation-consistency and reference-overlap positive control, not as independent proof of biological correctness.

In contrast, UROPA, ChIPseeker, and HOMER provide partially independent annotation semantics. Their disagreements with PeakLocus expose differences in query configuration, gene-assignment strategy, feature granularity, and native output vocabulary. The combination of a direct-overlap control and semantically distinct external tools therefore provides a more informative benchmark than any single comparator alone.

### Reference annotation file is a major source of cross-species variation

The species-stratified controlled benchmark demonstrates that exact label recovery depends on the annotation vocabulary encoded by the reference. Mouse and human GENCODE references use generic UTR features that can be resolved using CDS bounds and transcript strand, whereas FlyBase uses distinct feature naming and transcript conventions. PeakLocus showed essentially complete exact recovery in mouse and human but substantially lower exact UTR recovery in fly. This should not be read as evidence that the corresponding Drosophila loci are biologically non-exonic; rather, it reflects the sensitivity of an exact-label benchmark to the feature vocabulary and semantics of the supplied reference.

This observation has practical implications beyond PeakLocus. Software designed around one annotation ecosystem may require explicit normalization when used with another. For reproducibility, methods sections should report not only genome assembly and annotation release but also how feature names and transcript structures were interpreted.

### Start/stop-codon boundary semantics should remain explicit

A small number of mouse and human controlled discrepancies occurred at coding/UTR boundaries. PeakLocus currently treats start– and stop-codon records according to its transcript-boundary annotation convention rather than automatically calling those features CDS. Because the controlled masks were generated from five broad classes and did not encode start/stop codons as a separate truth category, such cases can be scored as disagreement even when the genomic placement is internally consistent with PeakLocus’s design. We therefore retain the current PeakLocus semantics and interpret these cases as boundary-definition differences rather than modifying the software to maximize benchmark score.

### Strengths of the benchmark design

The study combines two benchmarking regimes with complementary strengths. The real-world analysis evaluates broad and heterogeneous experimental peaks, multiple transcript isoforms, duplicate coordinates, and species-specific reference complexity. The controlled benchmark removes much of this heterogeneity and provides a balanced number of intervals from each RNA class, allowing class-specific F1 scores and species-level recovery to be compared directly. Together, these analyses distinguish coverage, concordance, and exact-class recovery rather than reducing tool performance to a single metric.

### Limitations

Several limitations should be considered. First, the controlled truth set is derived from the same GTF references used by the tools and is therefore not independent biological truth. Second, the strict six-category harmonization intentionally penalizes tools that return broader genomic categories or use different semantics. Third, exact class recovery can be sensitive to species-specific GTF feature names and to boundary features such as start and stop codons. Fourth, the benchmark evaluates interval annotation, not whether a reference transcript model is biologically correct in the assayed tissue or cell type. And finally, gene-ID concordance compares reported assignments under different algorithms and should not be interpreted as a universal ranking of gene-assignment correctness.

Future work could extend the benchmark with a dedicated challenge set containing UTR/CDS junctions, start/stop codons, splice boundaries, overlapping genes, and isoform-dependent segments. Such a challenge set would complement the current primary benchmark by explicitly testing cases in which more than one biologically defensible annotation may exist.

Together, PeakLocus provides a portable and transcript-aware approach for annotating RNA-associated genomic intervals with a focused RNA-location vocabulary. Its design integrates coordinate normalization, automatic strand handling, generic UTR interpretation, noncoding exon preservation, and transcript-aware intron derivation in a dependency-free Java command-line tool. Across 466,526 real experimental peaks, PeakLocus achieved 92.8% strict RNA-feature coverage and 99.5% concordance with a direct reference-overlap baseline among jointly annotated peaks. In a balanced 30,000-interval controlled benchmark, PeakLocus achieved near-complete exact five-class recovery in mouse and human, while cross-species and cross-tool differences highlighted the importance of reference vocabulary and annotation ontology. These results support PeakLocus as an interpretable RNA-centric annotation layer for RNA-modification and other transcript-associated genomic interval analyses.

## Software and Data Availability

PeakLocus source code and documentation: https://github.com/gsarfo-boateng/PeakLocus

Version evaluated in this manuscript: PeakLocus v1.0.0. The project is distributed under the MIT License.

Public benchmark datasets are available from NCBI GEO under GSE143821^18^, GSE192440 ^7^, and GSE114150 ^6^.

Benchmark scripts, harmonization code, controlled truth masks, and processed metrics should be deposited with the manuscript or released in the PeakLocus repository before submission [https://github.com/gsarfo-boateng/PeakLocus]

## Author Contribution

G.B.-S developed and implemented PeakLocus, designed and performed the benchmarking analyses, interpreted the results, and wrote the manuscript. S.P. contributed to the development of PeakLocus and reviewed the manuscript. S.S supervised the project.

## Ethical Statement

No new human or animal experiments were performed for this software study. All benchmark data were obtained from publicly available and their associated GEO numbers have been provided.

## Ethics Statement

No human or animal experiments we performed for the project. All benchmark data were obtained from publicly available sequencing datasets.

## Acknowledgements

We are thankful to the groups who generated and publicly released the sequence dataset used in this project.

